# Hippocampal Cdk5 is regulated by distinct stress paradigms in male and female mice

**DOI:** 10.64898/2025.12.23.696096

**Authors:** Kiara L. Rodríguez-Acevedo, Emily Ng, Michael D. Murphy, Paola Eusebio-Severino, Kyle S. Czarnecki, Elizabeth A. Heller

## Abstract

Anxiety and mood disorders have been rising since the COVID-19 pandemic, marked by social isolation and stress. Women are disproportionately affected by stress-related disorders, yet the interaction between social isolation and stress remains understudied in females. Cyclin-dependent kinase 5 (Cdk5) is an atypical kinase that exhibits sex-specific regulation following various stressors but is studied almost exclusively in males. To complement these findings, we examined sex differences in mouse hippocampal Cdk5 mRNA and protein following short and long restraint stress with or without concomitant social isolation stress. While short restraint stress was sufficient to promote anxiety-like in both male and female mice, it did not alter Cdk5. Rather, hippocampal Cdk5 mRNA increased only after a longer restraint stress duration followed by social isolation, and only in male mice. Hippocampal Cdk5 protein also shifted from the cytosolic to the nuclear compartment only in male hippocampus. Surprisingly, repeated restraint stress promoted exploratory behavior in both group-housed and socially isolated male and female mice. Social isolation alone increased Cdk5 protein in females only. Finally, under all conditions, female mice were less immobile and more exploratory than male mice. Together, these results provide new insight into the sex-specific regulation of hippocampal Cdk5 by the combination of psychological and environmental stressors.

**Highlights:**

1. Short 10-minute restraint stress induced anxiety-like behavior in both sexes without altering hippocampal *Cdk5* expression.
2. Long 1-hour restraint stress with post-stress social isolation elicited male-specific hippocampal *Cdk5* transcription and Cdk5 cytosolic-to-nuclear protein redistribution.
3. Repeated restraint stress increased exploratory behavior in both sexes, suggesting altered stress responsiveness.
4. Social isolation interacted with repeated restraint to alter hippocampal Cdk5 protein levels selectively in females.

## 1. Introduction

Anxiety and mood disorders have been rising steadily since the COVID-19 pandemic (Substance Abuse and Mental Health Services Administration, 2023), which was marked by widespread social isolation and persistent anxiety worldwide (World Health Organization, 2022). Although the social isolation associated with COVID affected both men and women, depression is more prevalent in women (National Center for Health Statistics, 2025). However, preclinical depression research has historically been conducted only on male rodents (Beery & Zucker, 2010); (Shansky, 2015) (Kaluve, et al., 2022), causing a critical gap in our understanding of sex-specific responses to stress including social isolation.

Stress exposure leads to a stress phenotype, which can manifest as reduced exploratory behavior and reduced social interaction, among other behavioral changes (Atrooz, et al., 2021) (Beery & Kaufer, 2014). Preclinical models involve either acute or chronic stress-exposure paradigms, such as immobilization through physical restraint (Molina, et al., 2023) or chronic social isolation (Pereda-Pérez, et al., 2013) (Grigoryan, et al., 2022). The latter is especially relevant given that rodents, like humans, are highly social animals. In male rodents, chronic social isolation increases inflammation and carcinogenesis (Hermes, et al., 2009) (Verza, et al., 2021), synaptic remodeling (Wang, et al., 2020) and depressive-like behaviors (Lukkes, et al., 2009). However, the impact of restraint stress and social isolation on the female rodent remains poorly understood. In this study we investigated the impact of different durations of physical restraint stress, social isolation, and their interaction on behavioral and molecular outcomes in male and female mice.

Here, we focused on Cyclin-dependent kinase 5 (Cdk5), a cell-cycle independent cyclin that is enriched in neurons and known to regulate stress- and depression-like behavior (Pao & Tsai, 2021). Although Cdk5 was first identified over three decades ago (Meyerson & Harlow, 1994) research on its role in stress and learning has excluded female rodents. Cdk5 plays a central role in both physiological and pathological processes in the brain, including in response to stress (Pao & Tsai, 2021) (Kimura, et al., 2014). For example, one hour of restraint stress increases Cdk5 protein levels and activity in the hippocampus of male mice for up to 24 hours post-stress (Papadopoulou, et al., 2015). Yet the effects of restraint stressors on the *Cdk5* gene remain largely unexplored. Our prior study finds that epigenetic activation of *Cdk5* gene expression in the ventral striatum mediates both stress and drug exposure in male mice (Heller, et al., 2016). Interestingly, epigenetic activation of *Cdk5* gene expression in the mouse hippocampus reduces fear memory in female but not male mice (Sase, et al., 2019). The current study aimed to advance knowledge on sex-specific regulation of Cdk5 by acute, physical restraint stress and chronic, environmental stress via social isolation. We hypothesized that restraint stress sex-specifically regulates hippocampal Cdk5 mRNA and protein abundance. We further hypothesized that prior social isolation would increase sensitivity to restraint stress. To approach these hypotheses, we exposed male and female mice to various durations of single or repeated restraint stress and chronic social isolation and measured hippocampal Cdk5 mRNA and protein.

## 2. Materials and Methods

### 2.1 Animals and Habituation

8 to 14-week-old male and female C57BL/6J mice were purchased from The Jackson Laboratory and maintained under standard laboratory conditions (12 hours/12 hours light/dark cycle, lights on 7:00 AM, 23 °C, unrestricted food and water). Mice were housed in same-sex groups of 3-5 or single-housed for isolation experiments.

### 2.2 Short restraint

Mice were handled 3 days before restraint day. Mice were immobilized for 10 minutes (min) using a 50 mL conical tube (with small holes drilled throughout for adequate respiration). Control mice were left in their home cage. Open field test was conducted 2-3 hours (hr) after restraint.

### 2.3 Long restraint

Mice were handled 3 days before restraint day. Mice were immobilized for 1 hr using a 50 mL conical tube as previously. Mice were euthanized 0, 3 or 24 hr after restraint. Control mice were handled and then left isolated until their euthanasia timepoint.

### 2.4 Repeated long restraint and isolation

Mice were group- or single-housed for 28 days before restraint stress, and housing conditions were maintained through designated restraint periods and behavioral tests. For restraint stress, mice were individually placed into a 50 mL conical tube for approximately 1 hr per day at 8:00 AM according to their assigned restraint stress period (D1/D7/D14). Following the last session of restraint stress, mice were exposed to stress-phenotyping behavioral tests. A subset of mice from each housing condition served as home cage controls and were not given stress. The light-dark box test was performed 1 hr after the final restraint session followed by the open field test.

### 2.5 Light-Dark Box Test (LDT)

The light-dark box test (LDT) was conducted 1 hr following the last episode of restraint stress using a repurposed conditioned place preference box. The apparatus consisted of two chambers partitioned by a small opening. Under direct fluorescent light, the light chamber was open-aired while the dark chamber was covered with aluminum foil. A camera was also mounted above the apparatus to record and track movement of the mouse. Before each animal test, the apparatus was thoroughly cleaned. Each experimental mouse was placed in the open area at the start of the test, and the animals were able to roam freely in the apparatus. Mice were observed directly and continuously by an observer and recorded on camera for 10 minutes. Stress-phenotyping involved measuring exploratory behavior as the time spent in the light chamber. Data was quantified using ANY-maze video tracking software (version 7.46, Wood Dale, IL).

### 2.6 Open Field Test (OFT)

The open field test (OFT) was conducted 2 to 3 hr after the last episode of restraint stress on an open-aired plastic box (45 cm x 45 cm x 45 cm). A camera was mounted above the apparatus to record and track movement of the animals. Before each test, the apparatus was thoroughly cleaned. Each experimental mouse was placed in the center of the box in the dark and allowed to roam freely in the apparatus. Mice were recorded on camera for 10 min. Stress-phenotyping measured exploratory behaviors (time spent in open area, average speed in the open area, and distance traveled) quantified using ANY-maze video tracking software (version 7.46, Wood Dale, IL).

### 2.7 Hippocampal Dissection

Mice were euthanized immediately after their final restraint stress session and behavioral testing by cervical dislocation. The whole brain was immediately extracted and chilled for a few seconds in phosphate-buffered saline (PBS, pH 7.4) with cOmplete, EDTA-free Protease Inhibitor tablets (Roche, 4693159001, Manheim, Germany) on wet ice. The brain was then placed in a chilled slicer matrix, coronally sectioned at 1 mm increments, and slices containing dorsal and ventral hippocampus were placed in a sterile dish with chilled PBS + protease inhibitor. Unilateral 2 mm-diameter biopsy punches for dorsal and ventral hippocampus (2 punches per region per mouse) were combined and collected in individual microcentrifuge tubes (4 total punches per unilateral mouse hippocampus) and immediately flash frozen in a metal tube rack placed on dry ice. Samples were stored at −80 °C. One unilateral sample per mouse was designated for mRNA quantification while the other was designated for protein analysis.

### 2.8 RNA Isolation and cDNA Preparation

Hippocampal tissue punches were homogenized, lysed, and fractionated, then cytosolic RNA was silica column purified using the RNeasy Mini Kit (Qiagen, Hilden, Germany). RNA concentration and purity were assessed using a NanoDrop One spectrophotometer (ThermoFisher, Waltham, MA). First-strand cDNA synthesis was performed using the iScript cDNA Synthesis Kit (Bio-Rad Hercules, CA).

### 2.9 Primer Design and Quantitative PCR (qPCR)

Primers for the genes of interest (Cdk5, GAPDH) were designed using the IDT PrimerQuest Tool (Integrated DNA Technologies, Coralville, IA), following default parameters for qPCR primer design. Sequences were selected to span exon-exon junctions, when possible, to reduce the likelihood of genomic DNA amplification. The resulting primers were synthesized by IDT and resuspended to a stock concentration of 100 µM. Working stocks were prepared at 10 µM in nuclease-free water. Primer sequences used in this study were as follows:

Cdk5:

Forward: GCTGCCAGACTATAAGCCCTAC; Reverse: TGGGGGACAGAAGTCAGAGAA GAPDH:

Forward: AGGTCGGTGTGAACGGATTTG; Reverse: GTGAGACCAGTGTAGTTGAGGTCA

qPCR reactions were performed using SYBR Green master mix (ThermoFisher, Waltham, MA) according to the following master mix recipe per reaction (10 µL total volume per well): SYBR Green Master Mix: 5.0 µL; Forward Primer (10 µM): 0.25 µL; Reverse Primer (10 µM): 0.25 µL; Nuclease-free H_2_O: 2.5 µL; cDNA template: 2.0 µL.

Each sample was run in triplicate, and a separate master mix was prepared for each primer pair to ensure consistency. Reactions were set up in a 384-well plate and run using a QuantStudio 7 qPCR thermocycler (ThermoFisher, Waltham, MA): 40 cycles of 1) denaturation at 95°C for 15 seconds, 2) Annealing at 60 °C for 30 seconds, and 3) Elongation at 72 °C for 1 minute. Melt curve analysis was performed post-amplification to confirm the specificity of each qPCR product. Cdk5 was normalized to Gapdh expression, and the ΔCT method was used to evaluate relative fold change expression (Livak & Schmittgen, 2001).

### 2.10 Protein Extraction and Western Blotting

Total proteins were extracted from hippocampal tissue using Radioimmunoprecipitation Assay (RIPA) buffer supplemented with protease and phosphatase inhibitors. For compartment-specific protein extraction, a lysis buffer containing 10mM Tris HCl PH 8.0, 10 mM NaCl, 3mM MgCl2 and 0.5% NP-40 was used. Lysates were then fractionated by centrifuging at 1500 rcf for 5 mins at 4 °C. The supernatant was taken as the cytosolic fraction, and the pellet was then further lysed with RIPA to extract nuclear proteins. All protein concentrations were determined using the Bicinchoninic acid (BCA) Protein Assay Kit (Pierce, ThermoFisher, Rockford, IL). Equal amounts of protein (10–30 µg per lane) were separated by SDS-PAGE and transferred onto polyvinylidene difluoride (PVDF) membranes. Membranes were blocked for 1 hr at room temperature in 5% bovine serum albumin (BSA) prepared in Tris-buffered saline with Tween 20 (TBST). After blocking, membranes were incubated overnight at 4 °C with primary antibodies targeting Cdk5 (Cell Signaling #2506) and housekeeping controls (A-tubulin [Cell Signaling #2144], GAPDH [Cell Signaling #2118], B-actin [Santa Cruz #sc-517582] or Histone 3 [Cell Signaling # 9715]). After washing, membranes were incubated for 1 hr at room temperature with Horseradish Peroxidase (HRP)-conjugated anti-rabbit (Cell Signaling #7074) or anti-mouse (Cell Signaling #7076) secondary antibodies. Protein bands were visualized using enhanced chemiluminescence (ThermoFisher, Waltham, MA) and imaged on a Bio-Rad ChemiDoc system (Bio-Rad, Hercules, CA). Band intensity was quantified using ImageJ software (version (version 1.53, Bethesda, MD) and target protein expression was normalized to housekeeping protein band intensity.

### 2.11 Statistical Analysis

GraphPad Prism software (version 10.2.2, La Jolla, CA) was used to perform statistical analyses and generate figures. Data are expressed as the mean ± standard error of the mean (SEM). An analysis of the variance (ANOVA) was employed for each analysis, followed by a post-hoc test with an alpha of 0.05 to correct for multiple comparisons. The Dunnett post-hoc test was used to compare all groups against a control, and a Tukey post-hoc test was used to compare all restraint periods against each other. Three-way ANOVA was used to compare all three variables (e.g., sex, housing, and restraint period), with post-hoc comparisons following interaction effects. Two-way ANOVA was used to compare two variables (e.g., restraint and housing), with post-hoc comparisons following interaction effects. Comparisons were considered significant at p-values below 0.05 and trending between 0.05-0.10.

## 3. Results

### 3.1 Short restraint stress induces anxiety-like behavior in males and females

To examine sex-specific responses to acute stress, male and female mice were exposed to a single 10-minute restraint stressor and subsequently tested in the open field test (OFT), a standard assay for assessing anxiety-like phenotypes (Kraeuter, et al., 2018) (Fig. 1A). We found that stressed males and females spent less time in the open arena and more time in the periphery compared to their non-stressed (control) counterparts (Fig. 1B-C). This reduction suggests that 10-minute restraint stress is sufficient to induce an anxiety-like phenotype in both sexes. Total distance travelled was similar between groups (Fig. 1D), indicating that locomotor activity was unaffected by restraint. Furthermore, we found a sex-dependent, stress-independent difference in time immobile, namely, females spent less total time immobile than males regardless of restraint condition (Fig. 1E) and exhibited fewer total episodes of immobility (Fig. 1F). This pattern may indicate that females engage in more active behaviors or maintain higher mobility at baseline and following short restraint stress.

**Fig. 1:**
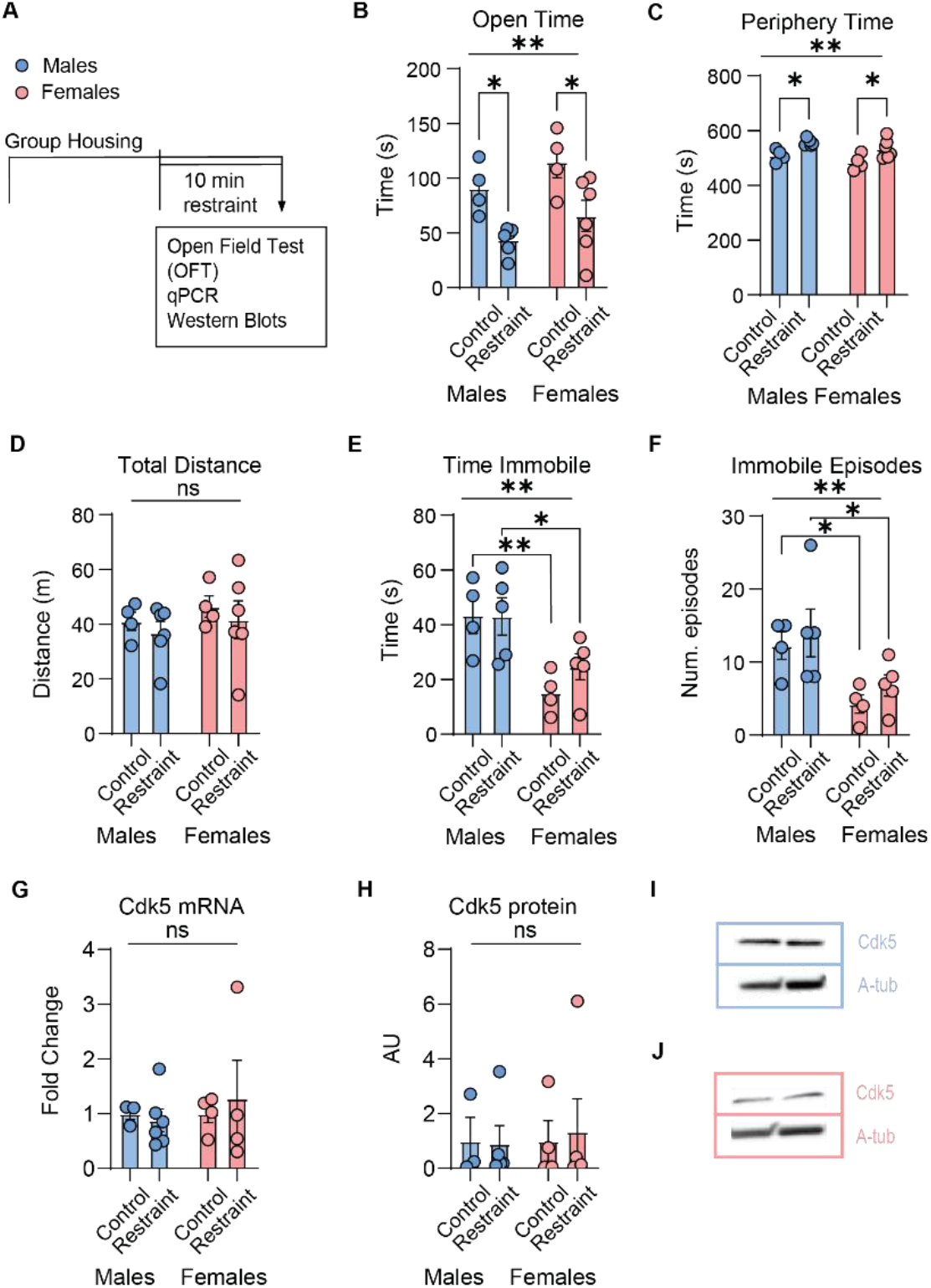
Short restraint stress inhibited open-field exploratory behavior in group-housed mice, with female mice showing less immobility than male mice. (A) Experimental design. A single 10-minute restraint stressor (B) decreased time spent in the open area during the open field test (OFT) in both sexes. Two-way ANOVA revealed a main effect of restraint (p = 0.0110) in both males and females (n = 4-5 per group), post-hoc analysis revealed an effect of control vs. restraint in both males and females (p = 0.0137 and p = 0.0103 respectively). (C) Time spent in the periphery was increased after short restraint in males and females. Two-way ANOVA revealed a main effect of restraint (p = 0.0110), post-hoc analysis revealed an effect of control vs. restraint in both males and females (p = 0.0137 and p = 0.0103 respectively). (D) Short restraint produced no changes in total distance traveled. (E) Females display reduced time spent immobile in both control and restraint conditions. Two-way ANOVA revealed a main effect of sex (p = 0.0013) on immobility, post-hoc analysis revealed an effect in females vs. males in both control and restraint conditions (p = 0.0057 and p = 0.0322 respectively). (F) Females also had fewer immobile episodes compared to males in both control and restraint groups. Two-Way ANOVA revealed effect of sex (p = 0.0048), post-hoc analysis revealed an effect in females vs. males in control and short restraint (p = 0.0330 and p = 0.0321 respectively). (G) *Cdk5* mRNA and (H) Cdk5 protein levels were unchanged by short restraint stress in (I) male and (J) female hippocampi. Data presented as mean ± SEM.

Given prior evidence that Cdk5 protein becomes activated after long restraint and chronic stress (Papadopoulou, et al., 2015), we examined hippocampal Cdk5 mRNA and protein, finding both unchanged after short restraint stress (Fig. 1H-I). These results suggest that 10-minute restraint stress is not sufficient to elicit detectable changes in hippocampal Cdk5 in male or female mice.

### 3.2 Long restraint stress elevates Cdk5 expression and alters protein dynamics in males only

We reasoned that *Cdk5* gene activation may require longer restraint stress duration. We thus employed a long, 1 hour, restraint paradigm that included post-stress isolation as an additional stressor. Social isolation during and after stress paradigms is common in the stress literature (Arakawa, 2018). We measured Cdk5 24-hr post-stress given evidence of increased Cdk5 protein and activity 3-24 hr post-stress (Papadopoulou, et al., 2015).

We found that hippocampal *Cdk5* mRNA levels were increased in male, but not female, 24-hr following 1 hr restraint and isolation (Fig. 2B-C) suggesting delayed transcriptional activation in males until 24-hr post stress. Total Cdk5 protein levels remained unchanged in both sexes (Fig. 2D, H). Cdk5 has been reported to exert some of its actions through activity-dependent nuclear translocation (Liang, et al., 2015). We thus performed cytoplasmic and nuclear fractionation to assess if we could detect activation of hippocampal Cdk5 protein in a specific cell compartment. We found that cytosolic Cdk5 protein was decreased in male, but not female, hippocampi at 24-hr following stress (Fig. 2E, I), and trending for increased nuclear Cdk5 protein in males only (Fig. 2F, J). Given this trend, we then measured the nuclear to cytosolic ratio of Cdk5 within animal. Indeed, males had increased nuclear to cytosolic Cdk5 (Fig. 2G) whereas females did not (Fig. 2K). In summary, long restraint stress combined with post-stress isolation led to a male-specific shift in cytosolic-to-nuclear Cdk5 ratio and an induction of *Cdk5* mRNA 24 hr post-stress.

**Fig. 2:**
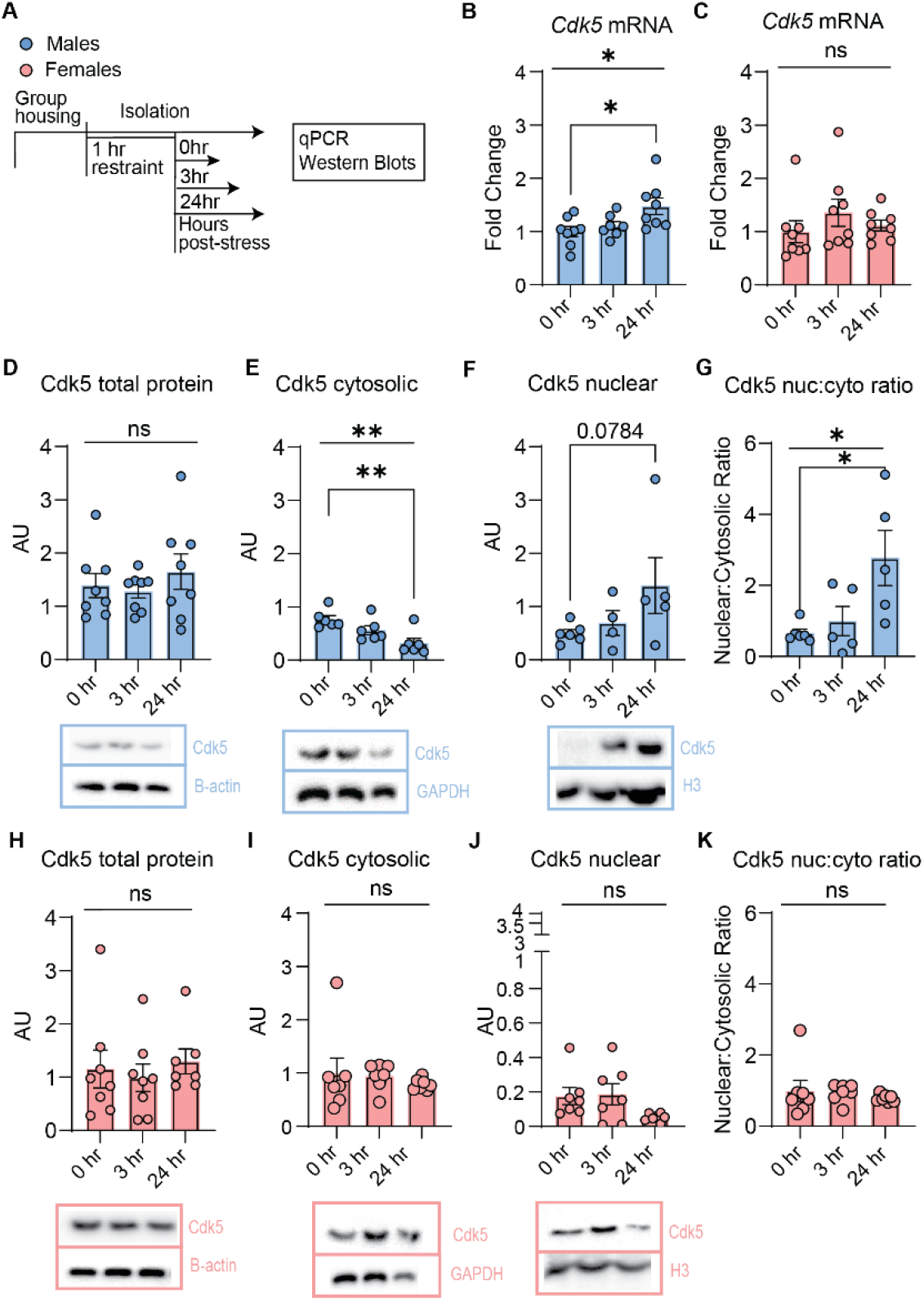
Long restraint stress increased Cdk5 mRNA and promoted Cdk5 nuclear import in male, but not female hippocampus. (A) Experimental design. A single 1-hr restraint session with post-stress social isolation (B) increased Cdk5 mRNA in male hippocampus 24 hr post-stress. One-way ANOVA revealed a main effect of post-stress time (p = 0.0208), post-hoc analysis revealed an effect of 0 h vs. 24 h post-stress (p = 0.0210) (n = 7-8 per group). (C) Female Cdk5 mRNA was unchanged. (D) There was no change in total Cdk5 protein in male hippocampus but (E) Cytosolic Cdk5 protein levels were decreased in male hippocampus. One-way ANOVA revealed a main effect of post-stress time (p = 0.0058), post-hoc analysis revealed an effect of 0 hr vs. 24 hr post-stress (p=0.0043). (F) Male nuclear Cdk5 protein levels were trending towards an increase in 0 hr vs. 24 hr post-stress (post-hoc analysis, p = 0.0784) (n = 5 per group). (G) One-way ANOVA revealed a main effect on the nuclear to cytosolic Cdk5 ratio (p=0.0176) with significant post-hoc in 0 hr vs. 24 hr post-stress (p=0.0187) in males. There were no apparent effects on female (H) Cdk5 total protein levels in the hippocampus or (I-K) compartment-specific Cdk5 protein. Data presented as mean ± SEM.

We next attempted to distinguish the regulation of Cdk5 by restraint stress from that of subsequent social isolation stress. We measured Cdk5 after long (1-2 hour) restraint stress at 0, 1, or 3-hr post-stress, during which mice were group housed (Fig. S2A). There was no change in *Cdk5* mRNA in males or female mice exposed to restraint stress compared to non-stressed controls (Fig. S2B-C). Cdk5 protein was greater in males after 1 hr restraint stress compared to 2 hr. However, we could not detect any differences of Cdk5 protein at any post-stress timepoint (Fig. 21D-E). Taken together, we found that hippocampal *Cdk5* increased only following the combination of long restraint stress and social isolation, in male mice only.

### 3.3 Long restraint stress promotes exploratory behavior in the open field test, despite reduced locomotion in males and females

Many human acute stress exposures are episodic in nature, rather than a single exposure (Weber, et al., 2022). We hypothesized that Cdk5 protein is sensitive to the combination of social isolation and repeated long restraint stress, but not short restraint stress alone. To test the effects of social isolation combined with repeated restraint stress, we subjected mice to long (1 hour) restraint for 1, 7, or 14 days (D1, D7, D14). We included social isolation as a variable, such that mice were group-housed or socially isolated 28 days prior to their restraint stress condition and during the 1-14 days of 1-hour restraint stress (Fig. 3A). We found a main effect of restraint on open field behavior in the OFT in both males and females. Surprisingly, one day of long restraint stress caused an increase in time spent in the center of the open field compared to non-stressed controls (Fig. 3B), regardless of housing condition (grouped or isolated). To validate whether this reflected an increase in exploratory behavior, we measured locomotor speed. Interestingly, we found that average speed in the open area was lower after one day of long restraint stress in both males and females regardless of housing condition (Fig. 3C).

**Fig. 3:**
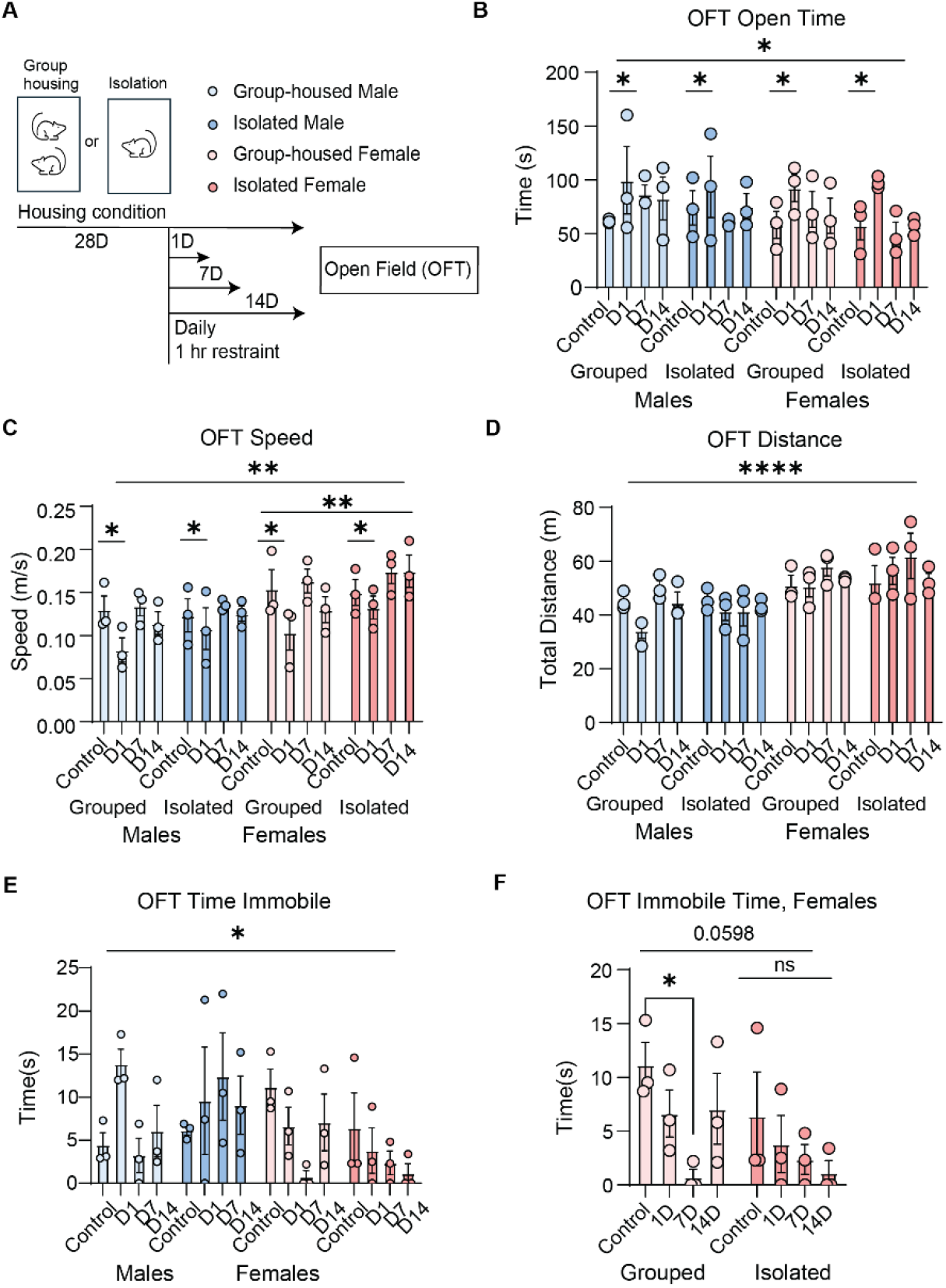
Long restraint stress increased time in the open field, despite reduced locomotion in both sexes. (A) Experimental design. Mice were group-housed or isolated and subjected to one (D1), seven (D7) or 14 (D14) days of long restraint stress. (B) Time spent in the open area during the OFT was increased after one day of long restraint stress in both sexes and housing conditions. Three-way ANOVA revealed a main effect of restraint (p = 0.0237), post-hoc analysis revealed an effect of Control vs. D1 (p = 0.0261) (n = 3 per group). (C) Average speed in the open area was reduced after one day of long restraint in both sexes. Three-way ANOVA revealed another main effect of restraint period (p = 0.0029), post-hoc analysis revealed an effect on control vs. D1 (p = 0.0320) and D1 vs. D7 (p = 0.019). Average speed in the open space was also generally greater in females than in males. Three-way ANOVA revealed a main effect of sex (p = 0.0011). (D) Total distance travelled increased in females independent of restraint day or housing. Three-way ANOVA revealed a main effect of sex (p < 0.0001). (E) Females spent less time immobile than males. Three-way ANOVA shows a main effect of sex (p = 0.0407). (F) When looking at the effect of housing condition in females, there was a reduction in immobility time after 7 days of restraint in group-housed females in comparison to group-housed controls (post-hoc analysis, p = 0.0363), with no effect in the isolated females. Data presented as mean ± SEM.

We also found a sex-specific effect on average speed in the open center, with females generally faster than males in all conditions. We also found that females traveled farther (Fig. 3D) and spent less time immobile (Fig. 3D) in the OFT compared to males, independent of housing condition (Fig. 3D). However, group-housed females showed reduced immobility at D7 of restraint, whereas isolated females did not (Fig. 3F). In summary, one-hour of restraint stress caused an increase in exploratory behavior along with a decrease in speed. We also found that females show increased locomotor activity and reduced immobility in comparison to males, with housing condition modulating some of these outcomes.

### 3.4 Repeated restraint increased time in light chamber of LDT in both sexes and social isolation increased Cdk5 expression in females

Given that one day of long restraint stress increased time spent in the open arena and reduced locomotor speed in the OFT in both males and females, we next examined anxiety-like and exploratory behavior using another behavioral test: the light-dark test (LDT). This test is well-validated in rodents to test for unconditioned anxiety-like behavior (Bourin & Hascoët, 2003) (Arrant, et al., 2013). We performed LDT in mice subjected to repeated long restraint stress (D1, D7, D14) with or without prior social isolation (Fig.3A). There were main effects of sex and restraint period on time spent in the light chamber. 7D of repeated restraint increased time spent in the light chamber of the LDT in both males and females with no effect of housing condition (Fig. 4B).

**Fig. 4:**
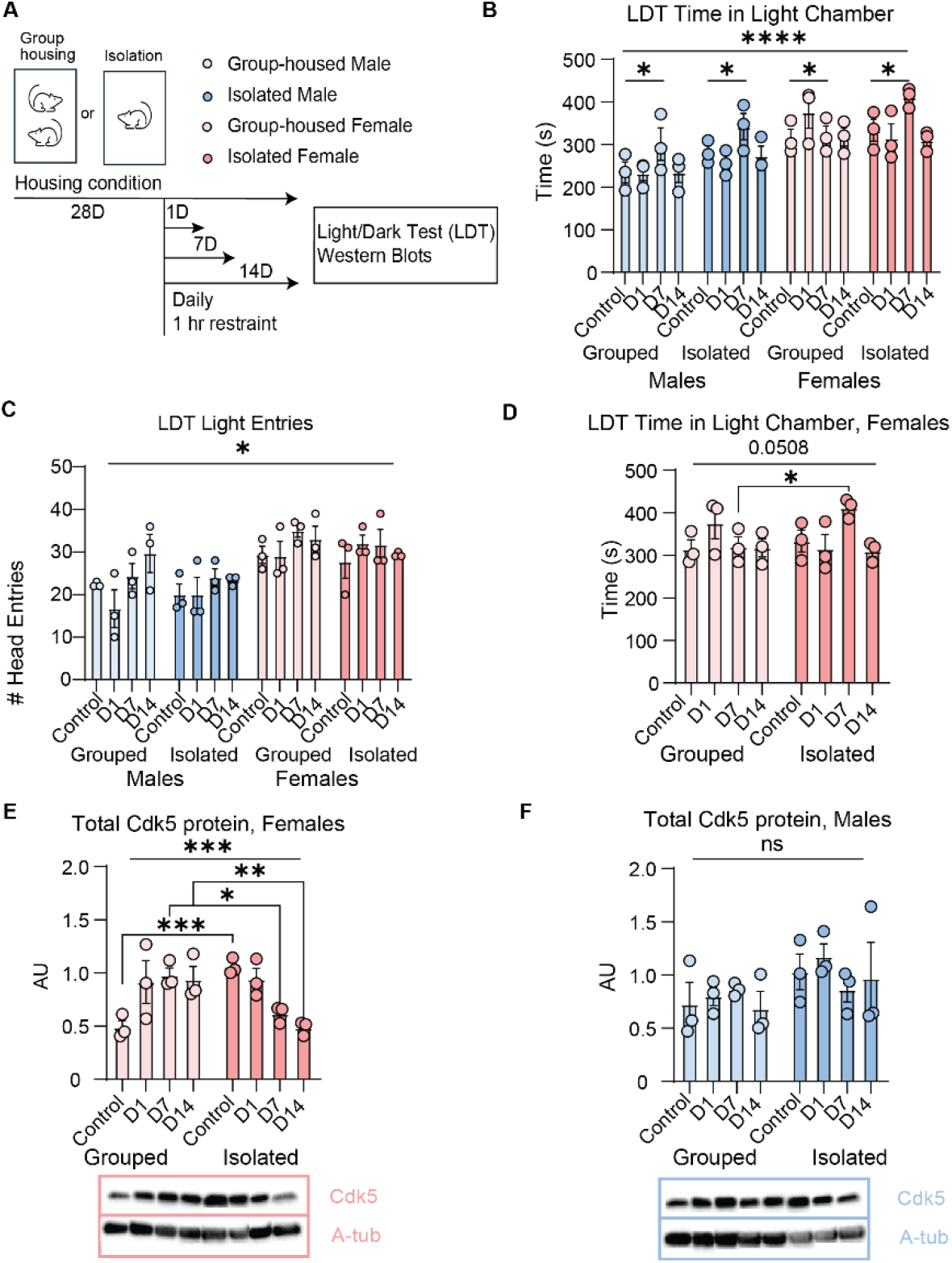
Repeated long restraint stress increased light time in LDT in both sexes and regulated Cdk5 protein in females only. (A) Experimental design. Priorly grouped-housed or isolated mice were subjected to one, seven, or 14 days (1D, 7D, 14D) of repeated long restraint stress. (B) 7D of repeated long restraint stress increased the time spent on the light side of the LDT test. Three-way ANOVA revealed a main effect of restraint (p = 0.0067), post-hoc analysis revealed an effect of Control vs. D7 (p = 0.0137, n = 3 per group). When looking at sex differences, females spent more time in the light chamber of the LDT than males, regardless of housing condition. Three-way ANOVA main effect of sex (p < 0.0001). (C) Females also made more entries into the light chamber than males, Two-way ANOVA revealed main effects of sex (p < 0.0001). (E) When probing for total Cdk5 protein levels by Western blot (n = 3 per group) in females, Two-way ANOVA revealed a main effect of the interaction between housing condition and restraint (p = 0.0004) with decreased Cdk5 in group-housed Controls vs. D1, D7 and D14 of restraint (p = 0.0370, p = 0.0151 and p = 0.03207 respectively) and increased Cdk5 in isolated Controls vs. D7 and D14 of restraint (p = 0.0395 and p = 0.0045 respectively). When looking at effects of housing in females, Cdk5 was increased in isolated vs. group-housed females in unrestrained controls (p = 0.0010). Conversely, there was a significant decrease of Cdk5 in isolated females at D7 and D14 of restraint (p = 0.0208 and p = 0.0053 respectively). (F) There were no changes in Cdk5 total protein in male hippocampus. Data presented as mean ± SEM.

When assessing head entries into the light chamber, there was a main effect of sex. Females made more light entries than males in all conditions (Fig. 4C). Examining housing conditions in females revealed that isolated females exhibited more light entries than group-housed females, particularly at D7 of restraint (Fig. 4D). These findings indicate interactions among sex, restraint, and housing in LDT performance.

We next assessed hippocampal Cdk5 protein levels. Cdk5 was affected by the interaction between housing condition and restraint in females (Fig. 4E) but not males (Fig. 4F). Specifically, hippocampal Cdk5 protein was increased in non-restrained, socially isolated females, and decreased in restrained females at D7 and D14. In summary, repeated restraint combined with isolation produces sex-specific behavioral and molecular effects: both sexes exhibit reduced avoidance of the light chamber after D7 of restraint regardless of housing condition while alterations in total hippocampal Cdk5 were shown in females only with effects by housing condition.

## 4. Discussion

### 4.1 Distinct behavioral responses to short versus long restraint in males and females

Animal model studies of acute and chronic stress have historically focused on males, leaving a significant gap in our understanding of female stress responses. By including both sexes across multiple stress paradigms, our study highlights how stress exposure produces sex-specific effects on behavior. A single exposure to short restraint elicited an anxiety-like response in the OFT in both sexes, manifesting as reduced exploratory behavior in the center open area. Alternatively, a single exposure to long restraint stress increased time spent in the center of the open field, although this behavior was accompanied by reduced locomotor speed, indicating that increased center time did not necessarily reflect enhanced exploratory drive. Locomotion is also impaired as measured by actophotometer after long restraint stress on day 1 but not on subsequent days in male mice (Chauhan, et al., 2015).

Seven daily exposures to long restraint stress produced an increase in time spent in the light chamber of the LDT, suggesting a shift from short stress reactivity toward altered coping strategies, including habituation. Notably, this increase in exploration was observed only after seven days of exposure to repeated restraint and was absent after 14 days, indicating that this behavioral adaptation may be transient. These findings are consistent with prior work in male rats showing habituation of stress hormone responses to repeated long restraint (Lovelock & Deak, 2018) (Márquez, et al., 2004) (Schmidt, et al., 2019) (Rabasa, et al., 2015). Together, these results underscore the importance of considering stress duration and number of exposures when interpreting behavioral outcomes in the OFT and LDT (Katz, et al., 1981) (D'Aquila, et al., 2000) (Chiba, et al., 2012). Females also exhibited reduced immobility in the OFT regardless of stress duration and condition, which may reflect a sexually dimorphic exploratory drive (Liiver, et al., 2023) or reduced behavioral inhibition that exists both in baseline conditions and after stress exposure.

### 4.2 The effects of social isolation alone are limited to females

Much literature exists on the effects of social isolation stress in rodents during adolescence (Fone & Porkess, 2008) (Bendersky, et al., 2021) (Burke, et al., 2017). In this study, we sought to distinguish the interaction and independent effects of *adult* social isolation and psychological stress in male and female mice. Interestingly, social isolation alone did not influence time spent in the open arena of the OFT in either sex. This is consistent with previous literature after both brief (Ferrara, et al., 2022) and prolonged isolation (Wang, et al., 2012) in adult male rats. Group-housed females exhibited less time immobile after 7 days of repeated restraint. This effect that was not observed in isolated females. This finding may point to a female-specific response to repeated stress that is attenuated with prior chronic social isolation stress.

At the molecular level, Cdk5 protein was increased in females after social isolation alone compared to group-housed controls. This suggests that, in females, Cdk5 is particularly sensitive to the experience of isolation itself. When isolation was combined with repeated restraint, this increase was no longer present, highlighting a distinct female-specific pattern of Cdk5 regulation under isolation conditions.

### 4.3 Sex-specific regulation of hippocampal Cdk5: nuclear shuttling in males vs. cumulative stress sensitivity in females

We previously found that the *Cdk5* gene is sex-specifically activated by classical fear conditioning (Sase, et al., 2019). Here we examined Cdk5 mRNA and protein in males and females after short and long restraint stress. Short restraint stress was insufficient to induce Cdk5 in male or female mouse hippocampus. This is consistent with previous work showing no change in Cdk5 protein levels after short duration restraint (30 minutes) in male rats (Adzic, et al., 2009) (Bignante, et al., 2010). In males, long restraint stress increased hippocampal *Cdk5* mRNA 24 hr post-stress, accompanied by a redistribution from the cytosolic to the nuclear compartment. Under homeostatic conditions, Cdk5 is anchored to the plasma membrane through its activator p35 (Asada, et al., 2012) (Sato, et al., 2007). Stress and other physiological or pathological stimuli can disrupt this membrane association, promoting Cdk5 nuclear translocation (Song, et al., 2016) (Liang, et al., 2015). Our compartment-specific western blots suggest that, in male mice, Cdk5 is regulated through this nuclear translocation mechanism. Cdk5 nuclear translocation may promote downstream signaling in the nucleus, including potential autoregulation of the *Cdk5* gene, which could influence behavioral adaptation to stress. Cdk5 interacts with epigenetic regulators such as HDACs and transcription factors including MEF2 (Fu, et al., 2013) (Gong, et al., 2003), making this mechanism plausible.

In contrast, female hippocampal Cdk5 levels were sensitive to the interaction between long repeated restraint and housing conditions: social isolation alone elevated Cdk5, whereas repeated restraint combined with social isolation decreased it. These findings indicate that, in females, Cdk5 protein upregulation is less rapidly initiated but more responsive to cumulative or interacting stressors, potentially contributing to sex-specific resilience. Sex differences in stress resilience have been described previously, although findings vary depending on the nature, duration, and timing of the stressor (Bangasser & Cuarenta, 2021) (Fallon, et al., 2020) (Kokras & Dalla, 2014) (Sood, et al., 2017). For example, females show greater resilience than males in modified social defeat stress (Pantoja-Urbán, et al., 2024) and learned helplessness paradigms (Shors, et al., 2007), yet results from the forced swim test remain mixed (Kokras & Dalla, 2014). Future work will be necessary to test whether sex differences in Cdk5 protein regulation translate into distinct transcriptional programs and behavioral phenotypes.

### 4.4 Implications for sex differences in stress resilience and vulnerability

Repeated stress exposure is a central feature of major depressive disorder, a condition with twice the prevalence in women compared to men (National Center for Health Statistics, 2025). To model this, we employed a repeated restraint stress paradigm while also considering housing conditions, given the frequent co-occurrence of psychological stress with social isolation in humans (Nguyen, et al., 2024). Behaviorally, both group-housed and isolated females showed lower immobility in the OFT following short and repeated long stress. Females also show greater preference for the light chamber in the LDT than males, consistent with enhanced exploratory drive and reduced anxiety-like behavior. Prior work in gonadectomized, estradiol-treated rats with or without repeated restraint stress demonstrated increased exploration in the OFT regardless of stress condition (Bowman, et al., 2002), supporting a role for estrogen in modulating exploratory and anxiety-related behavior.

A recently published study that isolated females exclusively in estrus and then observed OFT behaviors reported no effect of sex in the OFT (Smolensky, et al., 2024). Our study did not track estrous stage, therefore including females across all cycle stages, including high-estrogen phases. This broader sampling may help explain why we observed female-specific behavioral effects. Together, these findings suggest that female-specific mechanisms may contribute to altered stress responsiveness or coping strategies against certain stress-induced behavioral impairments. The female-specific sensitivity of hippocampal Cdk5 to cumulative stressors may help explain resilience under multiple stress exposure, potentially reflecting sex-specific molecular pathways relevant to stress-related disorders.

Overall, our findings expand the restraint stress literature by incorporating sex as a biological variable, examining the effects of multiple stress exposures, and identifying housing condition as a modulator of restraint stress reactivity in females. Importantly, we highlight distinct patterns of Cdk5 regulation in males and females, with Cdk5 protein nuclear translocation emerging as a potential male-specific adaptation, and cumulative stress sensitivity emerging as a key factor in females. These results emphasize the necessity of including females in preclinical stress research and suggest that sex-specific molecular adaptations, such as differential Cdk5 regulation, may contribute to the divergent vulnerabilities and resilience observed in mood and stress-related disorders.

## 5. Ethics Approval

All animal experiments were performed in accordance with the University of Pennsylvania's Institutional Animal Care and Use Committee and were conducted in accordance with the National Institute of Health Guide for the Care and Use of Laboratory Animals.

## 6. CRediT authorship contribution statement

KLR, EN, MM, and EAH conceptualized and designed the experiments.

KLR, EN, PES and MM conducted stress and behavioral paradigms.

KLR, MM, EN and KC carried out animal euthanasia and tissue collection.

KLR and EN conducted molecular experiments, including protein analysis and qPCR.

KLR wrote the first draft of the manuscript.

KLR, EN and EAH revised the manuscript.

## 8. Declaration of Generative AI and AI-assisted technologies in the writing process

During the preparation of this work the authors used ChatGPT(OpenAI) in order to revise for grammatical errors. After using this tool/service, the author(s) reviewed and edited the content as needed and take full responsibility for the content of the published article.

