## Supplemental Figure 2 for "Hippocampal Cdk5 is regulated by distinct stress paradigms in male and female mice"

**Supplemental Figures**

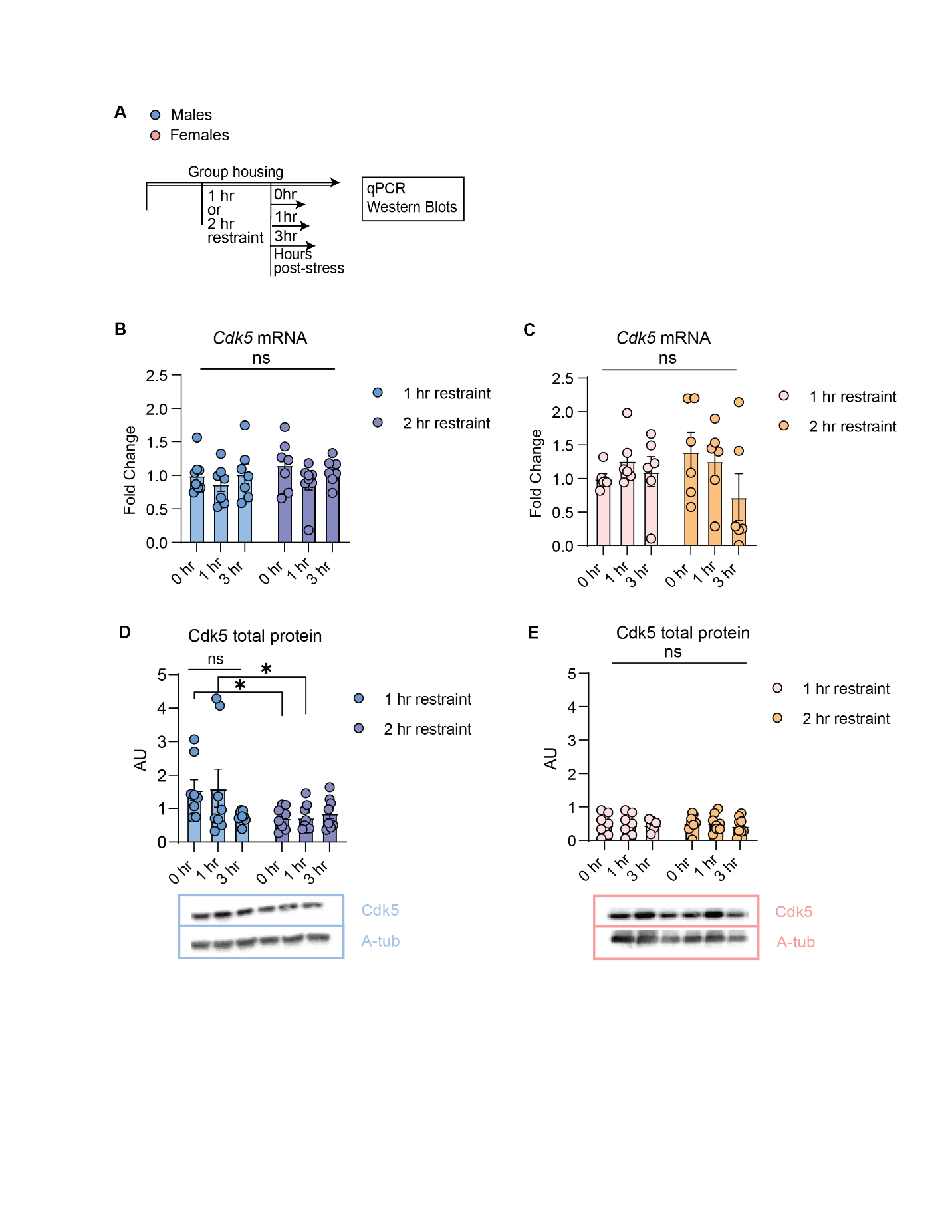

**Fig. S2: Long restraint stress in group-housed mice does not affect *Cdk5*.** (A) Experimental design. A single 1-hr or 2-hr long restraint session did not induce changes in *Cdk5* mRNA in either (B) male or (C) female hippocampus after 0-3 hr post-stress. Cdk5 total protein in the hippocampus was lower after 2-hr restraint stress in comparison to 1-hr restraint at 0 and 1 hr post-stress. Two-way ANOVA revealed a main effect of restraint stress duration (p = 0.0251) with post-hoc analysis significant at the 0 hr post-stress (p = 0.0419) and 1 h post-stress (p = 0.0350) timepoints. Data presented as mean ± SEM.
